# Pan-genomic and Polymorphic Driven Prediction of Antibiotic Resistance in *Elizabethkingia*

**DOI:** 10.1101/613877

**Authors:** Bryan Naidenov, Karyn Willyerd, Alexander Lim, Nathanial J Torres, William L. Johnson, Hong Jin Hwang, Peter Hoyt, John Gustafson, Charles Chen

**Affiliations:** 246 Noble Research Center, Department of Biochemistry and Molecular Biology, Oklahoma State University, Stillwater 74078, Oklahoma, USA; 4202 East Fowler Avenue ISA2015, Department of Cell Biology, Microbiology and Molecular Biology, University of South Florida, Tampa 33620, Florida, USA; 110F Henry Bellmon Research Center, Bioinformatics Graduate Certificate Program and Genomics Core Facility, Oklahoma State University, Stillwater 74078, Oklahoma, USA

**Keywords:** Nanopore sequencing, *Elizabethkingia*, Antimicrobial Resistance, Machine Learning, AMR prediction

## Abstract

The *Elizabethkingia* are a genetically diverse genus of emerging pathogens that exhibit multidrug resistance to a range of common antibiotics. Two representative species, *Elizabethkingia bruuniana* and *Elizabethkingia meningoseptica*, were phenotypically tested to determine minimum inhibitory concentrations for five antibiotics. Ultra-long read sequencing with Oxford Nanopore Technologies and subsequent *de novo* assembly produced complete, gapless circular genomes for each strain. Alignment based annotation with Prokka identified 5,480 features in *E. bruuniana* and 5,203 features in *E. meningoseptica*, where none of these identified genes or gene combinations corresponded to observed phenotypic resistance values. Pan-genomic analysis, performed with an additional 19 *Elizabethkingia* strains, identified a core-genome size of 2,658,537 bp, 32 uniquely identifiable intrinsic chromosomal antibiotic resistance core-genes and 77 antibiotic resistance pan-genes. Using core-SNPs and pan-genes in combination with six machine learning algorithms, binary classification of clindamycin and vancomycin resistance achieved f1 scores of 0.94 and 0.84 respectively. Performance on the more challenging multiclass problem for fusidic acid, rifampin and ciprofloxacin resulted in f1 scores of 0.70, 0.75 and 0.54 respectively.

## Introduction

Emerging antimicrobial resistance (AMR) is a global crisis. A recent report has predicted that by 2050 antimicrobial resistance will lead to 10 million deaths annually and cost the world’s economy upwards of $100 trillion (O’Neill, 2016;Tacconelli and Magrini, 2017). Currently, chemical assays that determine the minimum inhibitory concentration (MIC) are standardly used as a diagnostic tool to quantify antimicrobial resistance levels for cultured bacterial strains (Andrews, 2001). MICs quantify how susceptible, or resistant, a cultured strain is to selected antimicrobial drugs by observing the visible growth of the bacterium under antibiotic stress (Jorgensen and Ferraro, 2009). These protocols, however, are time-consuming and the interpretation of susceptibility for many antimicrobial/pathogen combinations have not yet been standardized (Horne et al., 2013). Furthermore, these procedures rely on the successful growth of bacterial isolates, making them incompatible with ‘unculturable’ bacteria (Vartoukian et al., 2010). As a result, full spectrum AMR detection remains challenging (Chitsaz et al., 2011).

With the advancement of sequencing technologies, single-molecule sequencing platforms are now regularly accessible and can overcome some of the disadvantages of phenotypic methods for AMR detection (Didelot et al., 2012;Fricke and Rasko, 2014;Lim et al., 2018). Genetic determinants conferring AMR have been identified in a few large studies. For example, the AMR profiles for 501 *Staphylococcus aureus* isolates were predicted from whole-genome sequencing (WGS) annotation results which achieved an overall sensitivity and specificity of 97% and 99%, respectively (Gordon et al., 2014). Another study investigating 681 *Neisseria gonorrhoeae* isolates achieved an acceptable major error rate (93% accuracy within one MIC doubling dilution and 98% for two) by regressing MIC phenotypes on genetic mutations of known AMR genes (Eyre et al., 2017). While the results are comparable to routine antimicrobial testing, they rely on the prior knowledge of single-gene products that relate to observable AMR phenotypes.

The wide range of expressed AMR phenotypes in the bacterial domain suggests that the genetic basis for the evolution and transmission of AMR is driven by a complex interplay of several factors. These include the rate at which resistance genes and mutations arise, the level of resistance contributed by the acquired genetic variants, and the relative fitness of the resistant mutants while under selective pressure from the drugs (Sommer et al., 2017). In addition, there are a large number of intrinsic chromosomal genes found in many bacteria, such as the marRAB operon (Vinue et al., 2013), which are involved in complex epistatic interactions regarding antimicrobial resistance expression (Gambino et al., 1993;Martin and Rosner, 2002;Lim et al., 2018). This can further render single AMR gene-based interpretations ineffective for predicting resistance phenotypes (Vinue et al., 2013).

Machine learning (ML) has led the way in cutting-edge prediction accuracy for vision tasks (Voulodimos et al., 2018), time-series problems (Ahmed et al., 2010), and the clustering of high-dimensional data (Assent, 2012). These methods have seen success in genomics for gene-finding (Libbrecht and Noble, 2015), predicting the functional consequences of protein missense mutations (Shihab et al., 2013), and genetic structure discovery using Markov clustering (Kopelman et al., 2015). ML is capable of dealing with high-dimensional interactions and nonlinear relationships in data, and has shown promise in using SNPs for predicting phenotypes that have a complex genetic architecture (Ban et al., 2010) and using k-mer counts (Nguyen et al., 2018). Recognizing the multifactorial, vertical, and horizontal genetic basis of resistance, we propose that AMR can be predicted for multiple phenotypes of a diverse group of multidrug resistant *Elizabethkingia* species by exploiting the capacity of cloud-knowledge driven ML approaches.

The Gram-negative rod genus *Elizabethkingia* demonstrates resistance to β-lactams and related antimicrobials due to the presence of multiple chromosomally-located β-lactamases (Bellais et al., 2000;Gonzalez and Vila, 2012). The high genetic diversity of *Elizbethkingia* also contributes to its highly variable MIC values for a broad selection of antibiotics (Perrin et al., 2017). In 2015 – 2016 in the states of Wisconsin, Illinois and Michigan (USA), 63 patients were found to have been infected by *Elizabethkingia anophelis*, which expressed multidrug resistance (Perrin et al., 2017). In this clonal outbreak, the individual pathogen isolates exhibited significant ecological structure with an uncharacteristic mutational spectrum. The temporal and spatial distribution of this population suggested on-going adaptation of the outbreak strain, possibly owing to an accelerated nucleotide substitution rate (Perrin et al., 2017).

Utilizing antimicrobial susceptibility and genomic data of 21 *Elizabethkingia* strains, including two newly completed strains sequenced with third-generation Nanopore long-read sequencing, this study examined the ML-powered predictability of AMR profiles. An *Elizabethkingia* core genome, built from the described strains, was established; and to leverage biological cloud data, other *non-Elizabethkingia* isolates were included to construct core-SNPs and pan-gene presence/absence matrices, with which AMR predictability was evaluated for several ML algorithms. In this report, we detail the methods used to produce efficient AMR prediction for both binary and multiclass resistance profiles.

## Materials and Methods

### *Elizabethkingia spp*., culture and DNA extraction

Single colony isolates of *Elizabethkingia bruuniana* ATCC 33958 and *Elizabethkingia meningoseptica* KC1913 were grown overnight in LB broth (10g NaCl/L) at 37°C under constant agitation. These cultures were then used to extract genomic DNA utilizing QIAGEN Genomic-tips DNA purification kits (QIAGEN, Valencia, CA) according to the manufacturer’s protocol.

### Antibiotic resistance profile evaluation

Bacterial isolates were grown and maintained as described previously (Johnson et al., 2018). Minimum inhibitory and bactericidal concentrations (MICs/MBCs, respectively) for each strain were determined by broth macrodilution following standard CLSI guidelines (Clinical and Laboratory Standards Institute, 2018). Overnight cultures were diluted in Mueller-Hinton broth (MHB) to an optical density at 600 nm of 0.01 where upon 1 mL was transferred to 13 mm x 100 mm sterile screw capped tubes containing 1 mL of antimicrobial. These tubes were subsequently incubated for 24 hours in a stationary incubator at 37°C, and the MICs were determined as the antimicrobial concentration that inhibited visual growth. Minimum bactericidal concentrations were determined by plating 100 μL from each tube at and above the MIC onto drug-free Mueller-Hinton Agar (MHA) and incubating for 24 hr (37°C). The MBC was determined as the lowest antimicrobial concentration in which no visual colonies were observed.

### Library preparation

DNA libraries were prepared separately for each *Elizabethkingia* isolate following the procedures outlined for the SQK-LSK208 2D sequencing kit (Oxford Nanopore Technologies, United Kingdom) with the following protocol adjustments. A total of 1.5 μg of gDNA was sheared in g-tubes (Covaris) at 4200 RPM for a targeted fragment size of 20 kb. End-repair was performed following the manufacturer’s recommended protocol for Ultra II End-prep enzyme mix (NEB). Adapter ligation reaction incubations were increased to 15 minutes. All bead clean-ups used 0.4x AMPureXP beads (Beckman Coulter, Brea, CA) for additional size selection and elutions were performed at 37°C for 20 minutes. DNA concentration of the library was quantified using Quant-IT PicoGreen^®^ dsDNA Assay Kit (ThermoFisher Scientific), measured on Synergy H1, hybrid multi-mode microplate reader (BioTek). Final DNA library yields were above the recommended 200 ng.

### Single molecular real time sequencing

Two R9.4 flow cells were prepared for two corresponding MinIONs, each connected to a separate Windows PC using a USB 3.0 connection. MinKNOW GUI application 1.0.8.0 from Oxford Nanopore Technologies (ONT) was used to validate the MinION connection and to monitor basic hardware details, like the number of active pores within each flow cell during sequencing runs. Pore count validation was completed beforehand, with the Platform QC command in MinKNOW. Flow cell priming was done according to the protocols provided by ONT for MinION use.

In a microfuge tube, 37.5 μL of running buffer (RBF), 25.5 μL of library loading beads (LLB) and 12 μL of *Elizabethkingia* DNA library were mixed to produce one loading library. The loading library mixture was carefully prepared for each species separately, to prevent fragmentation. The R9.4 flow cells received 75 μL of loading library via the SpotON port.

The sequencing runs were administered through the MinKNOW application, where each separate run was digitally labeled and the NC_48Hr_Sequencing_Run_FLO-MIN105_SQK-LSK208.py option was used for 2D R9.4 chemistries. After running the sequencing script, the flow cells were allowed to sequence for 48 hours, during which *E. bruuniana* was reloaded at the 24-hour mark. *E. meningoseptica* KC1913 was only sequenced for 20 hours, after which the flow cell provided no further sequencing capacity due to the depletion of nanopores.

### Assembly and polishing of *Elizabethkingia* genomes

The sequencing output from MinKNOW exists as the ONT FAST5 format, and Albacore 1.3.25 (ONT) was used to base-call the sequencing data. This transcribes the signal-level data into FASTQ sequences embedded within FAST5 reads. Extraction of the FASTQ data was completed using poretools version 0.6.0 (Loman and Quinlan, 2014). ONT changed the way that 2D FAST5 files are parsed causing a parsing problem in a critical downstream polishing tool for 2D FAST5 reads. Because of this change, all 2D reads were converted to 1D reads for the remainder of this study. The 1D template-strands and complement-strands were extracted with poretools using the switch: –type fwd,rev.

Per-read quality filtering consisted of a multi-step procedure to maximize read length and read quality for assembly. The reads, in FASTQ format, were subjected to a quality filter pass with a minimum Phred score of 12 using PRINSEQ (Schmieder and Edwards, 2011) with the – min_qual_mean switch. Reads with a length of 1,000 bp or lower were also discarded with PRINSEQ’s −min_len switch.

*De novo* assembly of each organism’s reads was completed with Canu v1.5 (Koren et al., 2017), a Celera Assembler successor designed to generate high-quality assemblies from Nanopore or PacBio long-reads. Canu was chosen because it provides higher assembly sequence identity than competing long-read assemblers, such as miniasm (Koren et al., 2017). Minimum overlap length was 500 bp and suggested genome size was 3.8 Mb and 4.5 Mb for *Elizabethkingia meningoseptica* KC1913 (Matyi et al., 2013), and *Elizabethkingia bruuniana* ATCC 33958 (Matyi et al., 2015) respectively.

After producing the initial *de novo* assembly with Canu, Nanopolish v7.1 (Loman et al., 2015) was used to improve the overall assembly quality for each sequence using a hidden Markov model. All original base-called, signal-level reads were re-extracted to tag the reads with identifying information for Nanopolish; these tagged reads were then aligned to their respective assemblies using BWA-MEM 0.7.15 (Li, 2013) using the −x ont2d switch. This command switch reduces the initial seed lengths and uses a relaxed scoring matrix, which allows the effective mapping of ONT’s noisy reads to the reference assemblies without producing large-scale fragmentation. After alignment, the produced SAM file was converted into the corresponding binary format (BAM file) using samtools (Li et al., 2009) using the view command and the −sB switch. Nanopolish was run in parallel to produce the consensus sequence for each assembly. The segmented output FASTA files were concatenated to complete the polished consensus sequence.

Signal-level data was used to capture methylation information (Simpson et al., 2017) across the genome. Nanopolish’s trained hidden Markov model was used to detect and compute likelihoods for potential 5-methylcytosine sites in the polished genome (Simpson et al., 2017). The final polished consensus sequence was then compared with the original unpolished assemblies from Canu using the software Mauve and its progressiveMauve algorithm (Darling et al., 2010). Since Mauve is not only an effective multiple genome aligner but also a variant caller, it was used to produce SNP and insertion/deletion (indel) tables for comparative purposes.

Circos (Krzywinski et al., 2009) was used to visualize the assembly data. For both polished genomes, a circular histogram was plotted to represent the per-chunk GC content. Each bar in the histogram represented the mean GC content for a 2,000 bp chunk of the genome, scaled from a minimum of 20% to a maximum of 50% GC content. Additionally, a heatmap representing methylation density was rendered using circos. Only methylation sites with a log-likelihood ratio greater than 3.5 were included. Darker regions contain methylation sites with a higher likelihood than lighter regions, and each region is represented by a chunk size of 3,000 bp.

### Annotation, cloud knowledge and multiple sequence alignment

Each polished assembly was submitted to the online RAST service (Aziz et al., 2008;Overbeek et al., 2014;Brettin et al., 2015) for annotation. Default settings of Classic RAST were used with Release70 as the RAST FIGfam version. Nanopore assemblies commonly contain deletions at homopolymer regions, resulting in frameshifts in the downstream DNA sequence; thus, the frameshift correction option in RAST was used to achieve better annotation results. Additionally, the building of a functional, metabolic model was selected as one of the options in RAST.

Prokka (Seemann, 2014) annotation relies on several databases, including UniProt, to predict CDS features in DNA. To provide an annotation comparison with RAST results, BLAST+ was used first; then HMMER3 was used as a sensitive search to mark features that were not found in the initial step. This provided an annotation solution that can be compared with earlier results produced from RAST. Prokka version 1.12 was used to annotate *E. bruuniana* and *E. meningoseptica*, using default parameters.

Finally, a third approach using the precise HMMER3 (Eddy, 1998) model was used to identify protein domains from an AMR database, Resfams (Gibson et al., 2015). MetaGeneMark (Noguchi et al., 2006) was used to mark putative protein-coding regions in both genomes with the gmhmmp −m command. The hmmsearch program from the HMMER3 suite was used, along with the Resfams HMM database v1.2, to identify potential protein domains associated with AMR. Identification of AMR gene clusters was done by filtering the output for regions with four or more AMR genes that had at most three non-AMR genes between any given gene pair. These results were visualized with circos. Putative promoters were predicted using the convolution neural network model software CNNProm (Umarov and Solovyev, 2017).

The vast extent of sequence data and available annotation information provide exciting opportunities for advances in biomedical sciences. To benefit from biological data available on cloud services and to enhance downstream analyses, the assemblies of 19 other strains of *Elizabethkingia* were acquired from the NCBI. Each assembly corresponds to a strain for which a known antibiotic resistance profile exists (see Supplementary Table 1 & Antibiotic Resistance Profile Evaluation section). This group of the 19 assemblies, paired with the two Nanopore assemblies (21 *Elizabethkingia* total), contained the *Elizabethkingia* species *E. bruuniana, E. meningoseptica, Elizabethkingia miricola, Elizabethkingia occulta*, and *E. anophelis* for use in core-genome construction.

Several *non-Elizabethkingia* strain assemblies were also acquired from the NCBI along with matching MIC results for vancomycin, clindamycin, fusidic acid, ciprofloxacin, and rifampin (see Supplementary Table 1). Following this, a “group” was created for each antibiotic, where membership to this group is determined by having an observed MIC value for that antibiotic. In total, 12 assemblies for vancomycin resistance, 7 assemblies for clindamycin resistance, 4 assemblies for fusidic acid resistance, 7 assemblies for ciprofloxacin resistance and 8 for rifampin resistance were retrieved from the NCBI that were not of the *Elizabethkingia* genus (see Supplementary Table 4). Many of the additional strains had MIC data for only one type of antibiotic.

For each antibiotic studied (vancomycin, clindamycin, fusidic acid, ciprofloxacin, and rifampin), an “AMR group” was formed containing strains that had corresponding MICs for the corresponding antibiotic. This generates five groups, each with a different number of individuals (Table 2). Separately, statistics were generated for the core-genome of *Elizabethkingia* strains only.

To generate a core genome for each group, a multiple sequence alignment of the assemblies for the strains within that group was first completed. The progressiveMauve algorithm (Darling et al., 2010) in Mauve was used to create six different alignments of the assigned bacterial groups. progressiveMauve was used instead of the original Mauve alignment algorithm for better scaling with multiple taxa and an improved scoring approach that handles the highly divergent genomes of the *Elizabethkingia* genus (Darling et al., 2010) and other included species.

### Core genome construction and SNP determination

Mauve produces a tabular “backbone” file containing alignments for DNA regions that are conserved between subsets of the genomes. These data are represented in a table with a header that describes each genome. Each column name in the header indicates the assigned ordinal-based index of the genome and also specifies if the given coordinate has been aligned in a reverse complement alignment. To extract only the core genome, all rows that did not contain conserved regions across all genomes were removed, leaving only the regions shared by all strains (core-alignments).

SNPs were called from within Mauve, using default settings for each alignment group. This did not include insertions or deletions. These SNP positions matched the regions listed within the “backbone” file that was produced prior to SNP calling. The resulting Mauve SNPs output is a tabular format file containing a row for each SNP site and a column for each individual in the alignment.

### SNP and gene predictor variables and response variables for AMR classification

To prepare the input data for downstream predictive algorithms, a predictor **X** matrix was constructed by directly loading the data from the Mauve SNPs file. Genotypic information from SNPs was represented as the encoded additive value for all polymorphisms on that site. The smallest value (starting at zero) represented the major allele. For the minor alleles, the encoded value increases as the frequency of the allele at that polymorphic site decreases. Effectively, the first minor allele will be encoded as 1, and in the case of multi-allelic polymorphisms, the next most frequency allele will be encoded as 2. The matrix was then transposed to adhere to the traditional structure of an input **X** matrix, in which all rows represent individuals and columns contain additively encoded SNPs.

While SNPs often function as genetic predictors of phenotypic traits, models utilizing only core-SNPs do not consider the presence of genes that are out of the core genome. To include these genes in the models, a list for all genes that exist within the strains of each group (pan-genes) was generated. A second, separate **X** predictor matrix was constructed using a presence/absence value. Each column represents the presence or absence of one putative pan-gene with a unique UniProt ID. A value of 1 is given for strains that contain the gene, while a value of 0 is given when the gene is absent. We consider this gene-centric approach appropriate in cases when genes, and not SNPs, are the true source of AMR, as seen in the relevant Tn1546 transposons that carry the *vanA* gene cluster conferring vancomycin resistant phenotypes (Dutka-Malen et al., 1994) or *fusB*-mediated fusidic acid resistance (O’Neill and Chopra, 2006).

Each set of predictors (pan-genes and core-SNPs) was used in evaluating AMR prediction. A final hybrid method was also evaluated by appending both matrices (SNPs and genes) together to form a combined, third predictor **X** matrix.

In order to assess the predictability of AMR with gene/SNP predictors, strains were labeled either “resistant” or “susceptible” to a particular antibiotic. Members of the *Elizabethkingia* genus have no standardized breakpoints so other species breakpoints are used for reference. Therefore, MIC values denoting resistance/susceptibility to the antibiotics vancomycin and clindamycin were based on the CLSI 2018 standards for *Enterococcus spp*. and *Staphylococcus spp*., respectively. For vancomycin resistance, any strain with a MIC value less than or equal to 4 was considered susceptible (CLSI 2018 M100 *Enterococcus spp*.), with the remaining strains being resistant. For clindamycin, the same protocol was used with a susceptibility label applied to MIC values of 0.5 or less (CLSI 2018 M100 *Staphylococcus spp*.). Fusidic acid was not considered for binary classification, as nearly every strain exhibited resistance to fusidic acid based on the reference breakpoints for *Staphylococcus spp*.

In each of the two AMR groups, the susceptible or resistant values for all individuals were represented by a **Y** vector, where each row in the vector (**y**_*i*_) is the observed phenotypic value for the corresponding row in the **X** matrix (**X**_*i*_). In the **Y** vector, a value of 0 represents a susceptible strain, while a value of 1 represents a strain that is labeled as resistant, with each respective group being assigned its own **Y** vector for that AMR category.

Higher resolution AMR prediction is possible by training models to predict a multiclass “resistance level” instead of a binary resistant/susceptible label. This was accomplished by assigning each strain a resistance level based on their MIC score. Resistance levels are therefore represented categorically as a sliding scale of AMR for the purposes of classification. Raw MIC results for each phenotype were collapsed into these resistance levels (see Supplementary Table 3 for resistance level assignments). The selection of ranges for binning MIC values for each phenotype was determined to maximize the uniformity of the categorical phenotypic distribution within the classes. This alleviated issues of outliers while minimizing the impact of very small numbers of individuals for the AMR category. Categories were encoded additively (similar to the SNPs), with the lowest resistance level encoded “1”; each AMR resistance category increases by one to reflect increasing resistance levels until it reaches the maximum value of that phenotypic category.

### Naïve Bayes

Naïve Bayes is a generative model used here to capture the posterior probability of the AMR classification given the SNP/gene predictors. This algorithm produces probability distributions based on the observed frequencies of the input variables and classifies using a simplified Bayes Rule. Using the probability chain rule with the assumption of variable independence, Naïve Bayes multiplies the probabilities of each specific class of variable and calculates posterior probabilities.

When occasionally considering a variable type that has not been observed in the training set the technique can result in a final probability of zero and numerical instability. Laplace smoothing was used to provide a small, non-zero probability to the probability for these types of classes. This is controlled by a smoothing parameter, **α**, a small constant which is added to the total number of actual observations for each type of variable, as well as increasing the total number of observed values. **α** is a hyper-parameter that must be optimized, where smaller values of **α** contribute less smoothing. The **α** values tested were 0.000001, 0.0001, 0.1 and 1.0. Each algorithm is assessed by stratified cross-validation where the test set will contain a pre-determined representation of classes. A uniform prior was used. The multinomial Naïve Bayes was conducted to predict the AMR category for individual strains using the sklearn package in python (Pedregosa et al., 2011).

### Decision tree and random forest algorithms

The decision tree is a non-parametric algorithm that can be used for classification or regression. Unlike the Naïve Bayes model, decision trees allow modeling of variable interactions and perform well on samples that cannot be linearly separated. In this tree structure, each node splits up samples based on a determined variable rule. This can be a threshold for continuous variables or a categorical value for discrete and categorical variables. To effectively split samples based on variables, the Gini impurity metric measures how well a particular node split in the tree will separate samples based on output category. Here, this calculation is given by

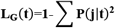

where ***t*** is the SNP or gene predictor and

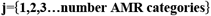

Minimization of the Gini impurity function maximizes the correct grouping of all samples during a node split.

A decision tree’s structure permits for excellent model interpretability and allows for the identification of important predictors. However, when the predictor count is much larger than the sample size, irrelevant SNPs/genes can produce trees that are fit on noisy, unrelated variables. When this ratio becomes particularly skewed, decision trees can become prone to overfitting and are inheritably sensitive to changes in training data. This lack of generalization occurs due to the large number of predictors, limited number of available samples and the high-variance nature of the SNP/gene predictors. In this assessment, all nodes were allowed to keep expanding until all leaves became pure and there was no maximum depth limit; sklearn was used (Pedregosa et al., 2011).

Random forests, an evolution of the traditional decision tree algorithm, has shown excellent modeling capacity by mitigating the issues of overfitting and high-variance nature of the decision tree (Breiman, 2001). This is done through an ensemble-based learning approach, similar to bootstrap aggregation, also known as “bagging” (Dietterich, 2000). Bagging follows the traditional bootstrapping of generating subsamples, with replacement, from the sample population and then allowing the model to classify based on those subsamples. This reduces the overall variance of the model while increasing the bias. Several models are then trained with different subsets and a majority vote is used when classifying test data. These random forests also only consider a subset of all the total predictors; subsetting by predictors provides similar advantages as subsetting the samples. As a result, these random forests often outperform stand-alone decision trees when the sample size is small and when the dataset is noisy (Dietterich, 2000), because they are fit to only a small subset of the samples of the variable. The sklearn package (Pedregosa et al., 2011) was used to classify bacterial strains into phenotypic categories, based on the predictors. The Gini impurity metric was used as in stand-alone decision trees with the sklearn package (Pedregosa et al., 2011).

Two hyper-parameters must be optimized to attain optimal performance, the number of decision trees in the ensemble and the number of variables to subset for each tree. To determine the ideal number of trees, tree counts of 5, 10, 20, 40, 60, 80, and 100 were tested. To identify the ideal number of SNPs/genes to subset, two subset counts were tested for each tree count:

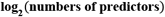

and

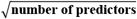

All nodes in the trees were expanded until all leaves became pure. Similar to earlier, there was no selected maximum depth limit.

### Boosting algorithm

“Boosting” algorithms work in a similar sense to “bagging” algorithms, in that several models are trained on a subset of the collected samples. Like random forests, subsamples are generated, with replacement, from the total sample population. However, unlike “bagging”, a modified sampling method is used. The probability of selecting any particular sample for training is increased or decreased depending on how well the models classify that subsample. Samples that are commonly misclassified are selected more frequently for training, and the opposite is true for correctly classified samples. This method attempts to minimize misclassifications of samples that are difficult to predict.

A “weak learner”, any classification algorithm that provides predictions that are only slightly better than random guessing, is applied to learn to classify the training subsamples (Freund and Schapire, 1997). Afterwards, all samples are associated with a weight value, which increases when they are classified incorrectly, and decreases when they are correctly classified. New weak learners are then generated, with the goal of minimizing the weighted error term (which is higher for misclassified samples) produced from the classification of new subsamples (Freund and Schapire, 1997). This process is then iterated until the weighted error term does not improve. This ensemble model of “weak learners” therefore emphasize the correct classification of “difficult-to-classify” samples. To correctly classify AMR categories, a decision tree was used as the “weak learner”, with a maximum leaf count of one; sometimes referred to as a “decision stump”, this one-level decision tree provides a simple classification model for data that is relatively unstructured, and is an effective base estimator for AdaBoost, the primary boosting algorithm used.

The primary hyper-parameter to optimize with AdaBoost is the number of decision stumps in the ensemble. Values of 10, 100, 500 and 1000 were tested. In cases where the number of predictors was less than 1000, the total number of trees in the ensemble was set to the maximum number of predictors in that dataset. The sklearn package (Pedregosa et al., 2011) was used with a learning rate of 1.

### k-Nearest Neighbor (k-NN)

k-Nearest Neighbor (k-NN) is a non-parametric algorithm that classifies test samples based on the Euclidean distance with training samples. These samples are represented in “feature space” as vectors, positioned in N-dimensional space, where N is the number of predictors for that AMR category. After populating the feature space with training data, new test data is classified based on the AMR category of the nearest samples (neighbors) in that space, using Euclidean distance as the metric, seen below (where ***p*** and ***q*** are the two feature vectors to compare):

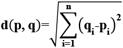

k is a user-defined constant, that determines how many neighboring samples are used in classification. When the nearest neighbors are of different categories, a majority vote is used to determine the class of the test sample. As such, k is usually odd to prevent balanced voting splits.

This algorithm is negatively affected when the number of predictors is very high (Beyer et al., 1999). As the total number of predictors increases, the distance to nearby neighbors approaches the distance to the most distant data point (Beyer et al., 1999). The model is particularly sensitive to its hyper-parameter k. Low values of k can overfit, while higher values can underfit. The sklearn python package (Pedregosa et al., 2011) was used to model the performance of k-NN with six different values for the k hyper-parameter being evaluated: 1, 3, 5, 7, 9 and 11.

### Support Vector Machines (SVMs)

Support vector machines (SVMs) were also evaluated, due to their effectiveness at classifying high-dimensional data like biomarkers and microarrays (Clarke et al., 2008). SVMs work by generating a N-dimensional hyperplane, so as to separate samples by their classification category. This hyperplane is defined as being an N-dimensional plane that sits between two margins (for binary decisions), and these margins are produced by the data points of opposing classes which are closest to the decision boundary. These data points of interest are designated “support vectors”. For data that is linearly classifiable, a linear SVM can be used; however, for non-linearly classifiable data, kernels are often employed to map these data into a feature space where a linear hyperplane can be constructed.

SVMs can use several strategies to evaluate more than two classes. In the case of classification as a multiclass problem, the “one-against-one” method trains a classifier for each different pair of categories for a total of

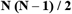

classifiers. At test time, all the classifiers are tested on the test samples, where each classification is a vote for that particular class. The class with the most votes is determined as the correct category for that sample.

A key hyper-parameter for SVMs is the constant **C**, which acts within the soft margin cost function; this **C** term controls the tightness of the two margins used to produce the hyperplane. Larger **C**’s will produce tighter margins, resulting in less misclassified training samples. A smaller **C** will produce larger margins, allowing for the misclassification of some training samples. Smaller values may help deal with outliers and form a more generalizable hyperplane by trading error penalty for model robustness. In this study, the penalty hyper-parameter **C** term was tested with the following values: 0.01, 0.1, 1.0, 10, 20 1000 and 10000, and the hyperplane was optimized as a dual optimization problem using the sklearn library (Pedregosa et al., 2011). Both linear SVMs and SVMs with radial basis function kernels were tested.

### Evaluation of prediction accuracy

To evaluate the performance of the predictive models, accuracy of the classification algorithms was determined with 18,000 iterations of stratified shuffle cross-validation and a computed f1 micro-score, representing the harmonic mean of both recall and precision. The f1 micro-score is more descriptive than calculating classification accuracy as a percentage. Stratified shuffling was used in combination with cross-validation so as to maximize the uniformity of the category distribution in the test and training sets.

The test set sample size for binary classification of AMR for vancomycin and clindamycin was 6, which was performed to allow a reasonable number of test samples to be involved in classification assessment. Multiclass “resistance level” classifications were evaluated for all five AMR types, with a test set size of six and the same number of iterations described above.

## Results

### Antibiotic minimum inhibitory and bactericidal concentrations

MICs and MBCs, for all antibiotics investigated, varied among the *Elizabethkingia* species and strains investigated. The ciprofloxacin MICs ranged from 0.125 mg/L to 1 mg/L and MBCs ranged from 0.5 mg/L to 2 mg/L as shown in Supplementary Table 1. The clindamycin MICs ranged from 0.0625 mg/L to 1 mg/L and MBCs ranged from 0.0625 mg/L to 8 mg/L (Supplementary Table 1). The rifampin MICs ranged from 0.0625 mg/L to 1 mg/L and MBCs ranged from 2 mg/L to 32 mg/L. The fusidic acid MICs ranged from 4 mg/L to 128 mg/L and MBCs ranged from 4 mg/L to 256 mg/L. The vancomycin MICs ranged from 2 mg/L to 64 mg/L and MBCs ranged from 4 mg/L to 64 mg/L.

### Nanopore R9.4 sequencing yield of *E. bruuniana* and *E. meningoseptica*

The original 2D sequencing of *E. bruuniana* ATCC 33958 yielded 212,265 total reads (Table 1). During sequencing, low-quality reads or reads containing errors were filtered out, leaving a total of 166,167 quality reads. The errors in base-calling were attributed to software exceptions (1,392 reads) and failed 2D software base-calling (33,981 reads), while 10,725 reads did not pass the base-caller’s built-in quality filter. Throughout sequencing, the mean Phred score distribution of 2D reads was maintained at 16 at any given hour, except during reloading, where the quality distribution fell to 14. The median 2D read length was 5.78 kb and the average 2D base-calling accuracy was 0.92. Template and complementary base-calling accuracy was 0.85 and 0.80, respectively.

**Table 1.** Comparison of Nanopore R9.4 2D sequencing statistics. All values reported here are from reads that passed the quality threshold.

|  | *E. bruuniana*<br>ATCC 33958 | *E. meningoseptica*<br>KC1913 |
| --- | --- | --- |
| Total quality yield (megabases) | 1,090 | 330 |
| Total number of quality reads | 166,167 | 45,786 |
| Median read length (kilobases) | 5.78 | 6.53 |
| 2D Median quality score (phred score) | 15.8 | 16.3 |
| Template median quality score (phred score) | 8.4 | 8.8 |
| Longest read (kilobases) | 32.8 | 42.4 |

Sequencing *E. meningoseptica* KC1913 with Nanopore long-reads produced a total of 57,521 2D reads. Of these reads 1,190 software exceptions, 8,111 instances where base-calling failed, and 2,434 reads that didn’t pass the quality filter were removed, yielding 45,785 passed reads. The median 2D read length was 6.53 kb. The mean quality score was 16 for the entirety of the sequencing run. Average 2D base-calling accuracy was 0.93, while template and complementary base-calling accuracy was 0.86 and 0.81, respectively.

During 2D sequencing of *E. bruuniana*, per-hour base-pair yield peaked at 2 hours (54 mb). A second peak of 46 mb of DNA sequenced was observed during reloading at 24 hr, followed again by a rapid decrease, converging to nearly zero bases sequenced per hour by 45 hr. *E. meningoseptica* was sequenced for a total of 20 hours (due to hardware failure), during which it received no library reloads. The highest hourly yield peaks (30 mb) occurred during hours 1 and 3, and yield rapidly diminished to 3 mb per-hour by hour 20.

Following the results of read filtering, Figure 1 shows the read length distributions and quality score distributions for both sequenced strains. *E. bruuniana* and *E. meningoseptica* has median 2D quality scores of 15.8 and 16.3 respectively (Table 1).

**Figure 1.**
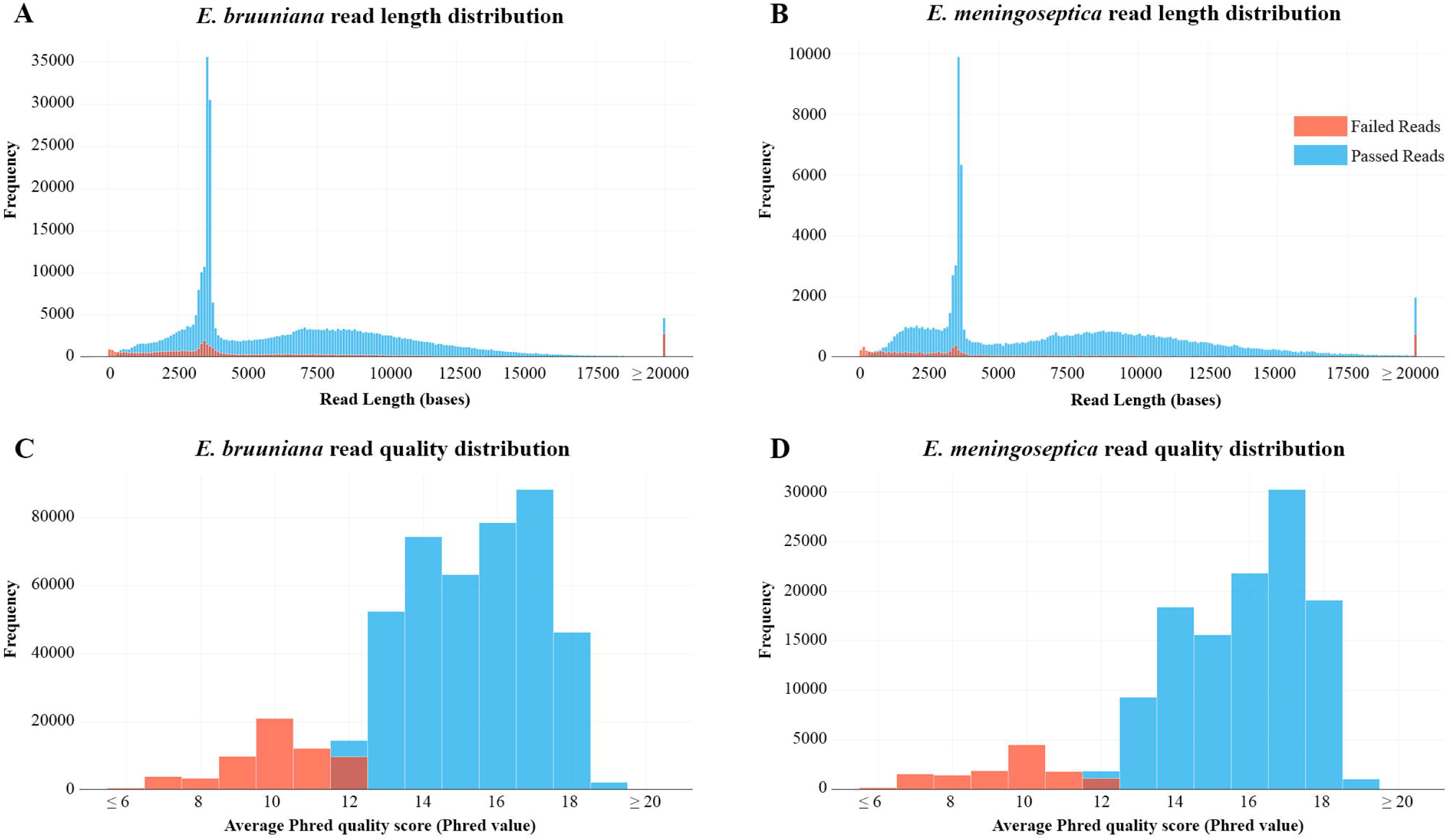
A.) Read length distribution after extraction and filtering of raw read data of *Elizabethkingia bruuniana* B.) Read length distribution after extraction and filtering of raw read data of *Elizabethkingia meningoseptica* C.) Per-read quality distribution of *E. bruuniana* D.) Per-read quality distribution of *E. meningoseptica*

### Genome assemblies and polishing

*De novo* assembly of *E. bruuniana* and *E. meningoseptica* resulted in a single contig for each genome. The assembly of *E. bruuniana* (ATCC 33958) was 4,626,295 bp long, with a mean GC content of 35.9% and an average read coverage of 610x. The assembly of *E. meningoseptica* (KC1913) was 3,862,237 bp long, with a mean GC content of 36.5% and an average read coverage of 220x.

After polishing, the assembly size of *E. meningoseptica* increased slightly to 3,889,109 bp, primarily due to corrections in deletion errors at homopolymer regions. Deletions were the most common per-base assembly error, consisting of a total of 21,321 gap regions for *E. meningoseptica*. These occurred almost exclusively at homopolymer repeat regions. The majority of gap regions contained only a single deletion (80% of all gaps were single-base deletions). A much smaller quantity of insertion errors (74 regions) that met the reporting criteria were also corrected by the polishing process. All of these reported regions contained only a single-base insertion error. Substitution errors were also minimal; only 1,273 substitutions were corrected within the *E. meningoseptica* assembly, and the majority of these substitutions (> 46% of all substitutions) were made up of G to A and C to T corrections.

Similar polishing results were also achieved with *E. bruuniana*, where deletions were again the largest source of corrected errors (20,133 deletions), resulting in a slightly larger genome sequence (4,651,278 bp). A total of 279 insertion errors and 990 substitution errors were corrected. Unlike in *E. meningoseptica*, no particular substitution error dominated the corrections.

The assemblies of the two newly assembled *Elizabethkingia* genomes have been deposited in a NCBI BioProject (for early access, contact).

### Genome annotation, AMR gene identification and methylation prediction

Annotation with RAST discovered 5,114 putative features in *E. bruuniana* ATCC 33958 and 4,887 putative features in *E. meningoseptica* KC1913. In *E. bruuniana* we found that RAST had putatively identified 19 ß-lactamases and 18 efflux pumps. In *E. meningoseptica* there were 14 putatively identified ß-lactamases and 19 efflux pumps.

Annotation with Prokka produced 5,480 features in the *E. bruuniana* genome, among which 5,426 were identified to be coding regions, 53 identified as tRNA genes and one feature identified as a tmRNA gene. In *E. meningoseptica*, there were 5,203 features found in total, 5,152 of these features matched coding regions, 50 matched tRNAs, one matched tmRNA and one was identified as a repeat region. Details of Prokka annotation of *E. bruuniana* genome can be seen in the Supplementary Table 5, and Supplementary Table 6 for *E. meningoseptica*.

Identification of putative AMR genes thought be related to vancomycin resistance, using Prokka, found that all *Elizabethkingia* strains contained a *vanW* gene, with *E. meningoseptica* KC1913 additionally containing a *vanB* gene (Supplementary Table 2). Several *erm* genes were found associated with clindamycin resistance; however, only two isolates had observed clindamycin MIC values to compare with: the gut Gram-negative pathogen *Parabacteroides goldsteinii* 910340 and Gram-postive pathogen *Enterococcus faecium* 805447/07. Both isolates exhibited strong clindamycin resistance based on the observed MIC values of 256 and 8 mg/L respectively (Supplementary Table 1). No gene annotations were found relating to fusidic acid, rifampin and ciprofloxacin resistance in any of the species investigated.

High-quality annotation with the Resfams database and HMMER3 identified 569 unique putative genes associated with AMR in *E. meningoseptica* KC1913 and 685 in *E. bruuniana* ATCC 33958. The 15 gene clusters in KC1913 and the 12 clusters in ATCC 33958 are visualized in Figure 2.

**Figure 2.**
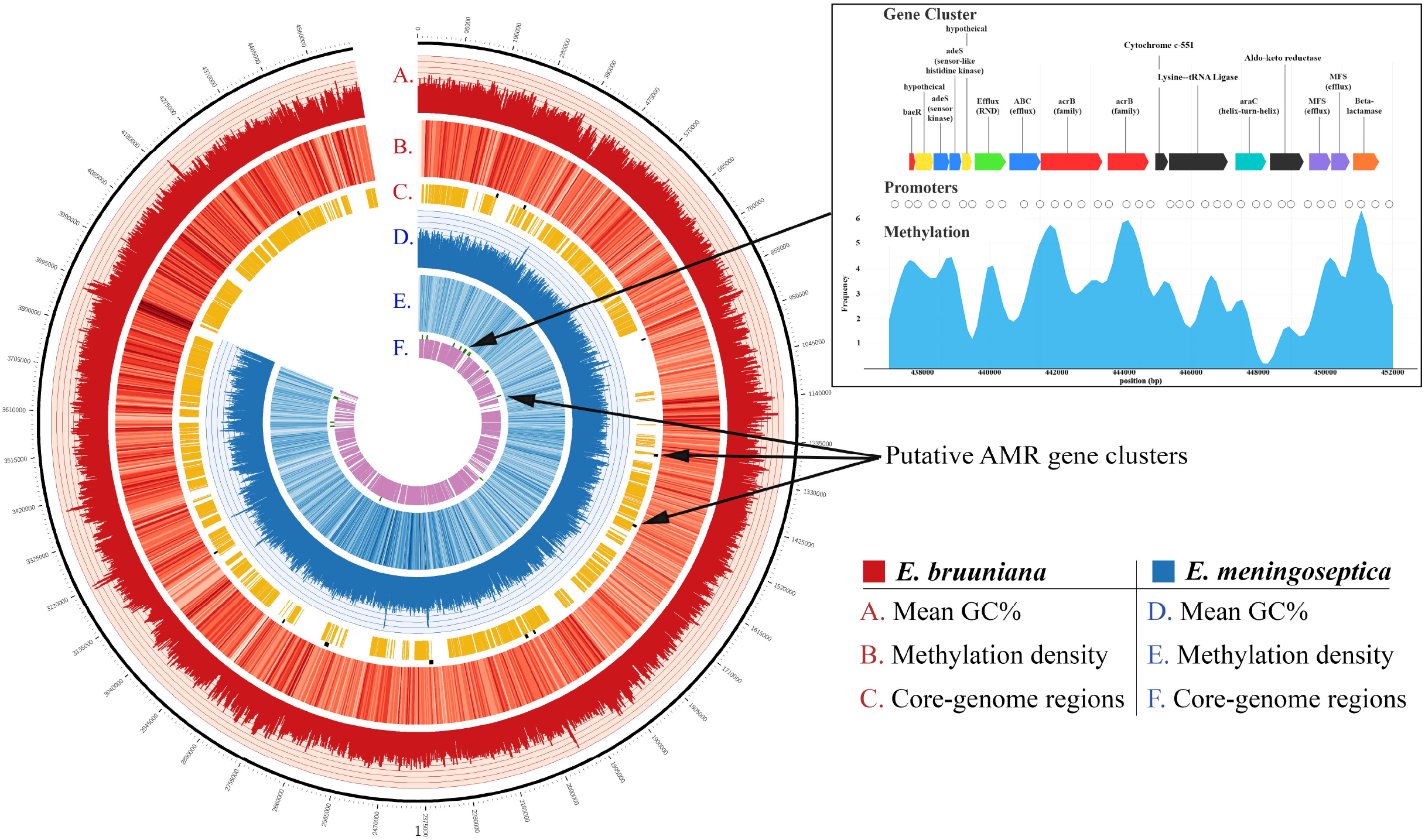
Competed genomes of both *E. bruuniana* (red) and *E. meningoseptica* (blue) are displayed with histograms representing average GC content (ranged 20% − 50%) in the outermost circles of each genome. The middle genome circles display a heatmap indicating methylation frequency (darker regions indicate high methylation frequency). The inner genome circles indicate conserved regions belonging to the *Elizabethkingia* core genome found in each respective genome. Putative AMR gene clusters, identified by HMMR3, are shown on the outer edges of the core genome circles.

Prediction of methylated cytosine sites using Nanopolish generated 82,175 non-redundant sites of potential cytosine methylation for *E. meningoseptica*. Methylation was predicted in 29,545 of the total reads and totaled 5,566,105 sites, many of them shared across separate reads. Similar results were found with *E. bruuniana*; 89,788 non-redundant cytosine methylation sites were predicted. In total, 17,316,177 shared sites were found over 109,339 reads. The distribution of methylation is shown as a heatmap in Figure 2. Dark blue regions (for *E. meningoseptica*) and dark red regions (for *E. bruuniana*) show sites with high frequencies of predicted cytosine methylation regions. In *E. bruuniana*, methylation-rich CpG-rich regions often appeared in non-core genomic regions of the strain (Figure 2B and 3C).

**Figure 3.**
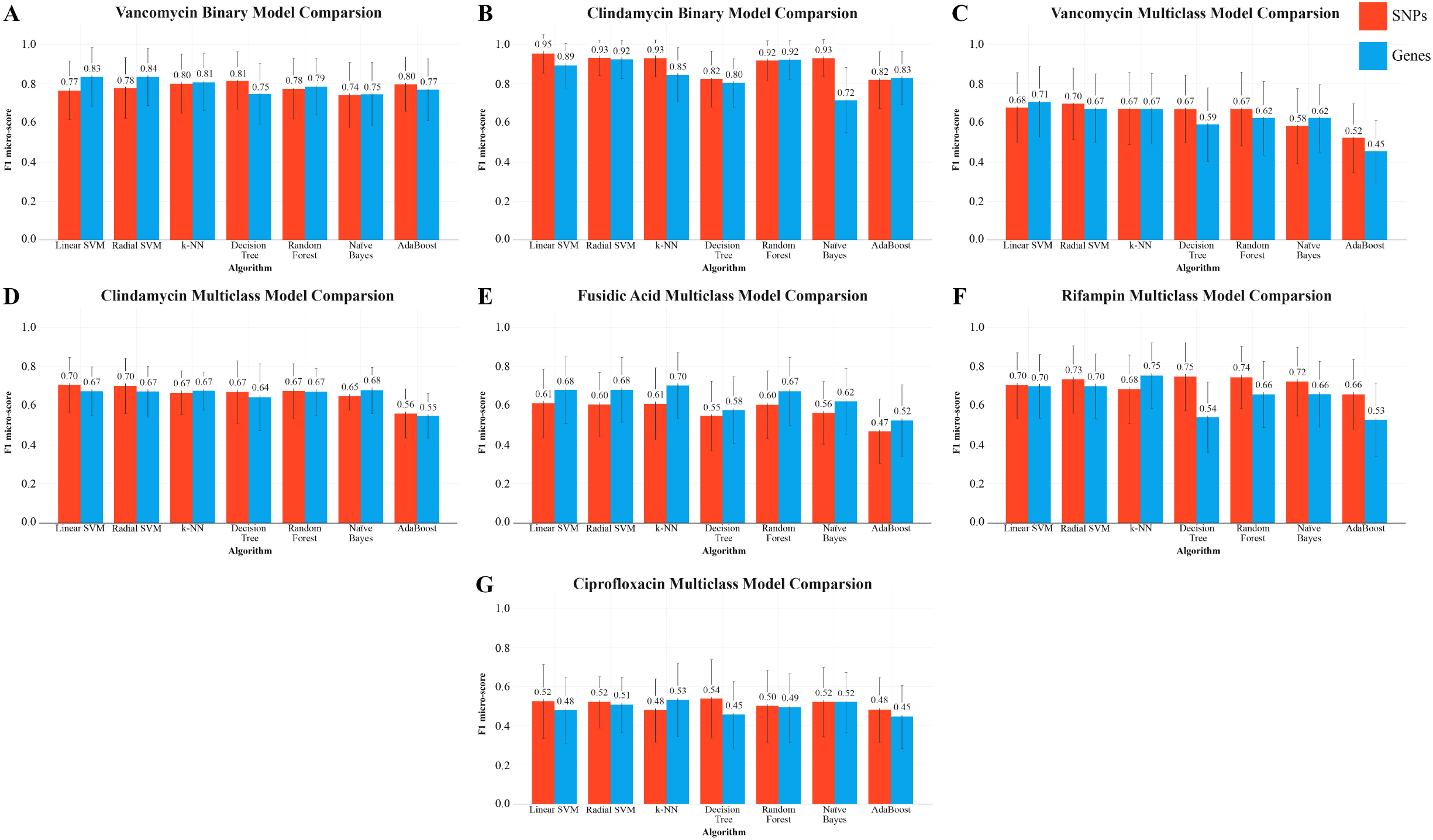
f1-micro scores for each algorithm (with their best respective hyper-parameters), for each AMR group. Mean f1-micro score over the given iterations for that group is shown by the values above the bars and standard deviation is shown by the error bars. The binary prediction algorithms are show in panels A. and B. The multiclass classification is shown in D. – G.

### Summary of core and accessory genomes, and SNP determination

Multiple-sequence alignment of the *Elizabethkingia*-only genomes produced a larger core-genome size when compared to the other groups (Table 2), resulting in core genome size of 2,658,537 bp, with 32 core-AMR genes and 77 pan-AMR genes based on Uniprot IDs. It also produced the largest number of called SNPs (712,703 SNPs). When including bacterial species from different taxonomic groups, the increased genetic diversity drastically reduced the size of core-genomes, as well as the number of SNPs called. However, the reduction in core-genome size was not proportional to the reduction of SNPs. The largest *non-Elizabethkingia* core-genome was formed from the genomes in the vancomycin MIC group, with the fusidic acid MIC group being only a few thousand bases smaller. The complete core-genome of the two sequenced isolates KC1913 and ATCC 33958 is visualized in yellow in Figure 2.

**Table 2.** Core genome and SNP statistics for each phenotypic group formed from participating strains.

| Phenotype | Number of strains | Core genome size (bases) | Number of SNPs |
| --- | --- | --- | --- |
| *Elizabethkingia*-only | 21 | 2,658,537 | 712,703 |
| Vancomycin | 33 | 27,066 | 11,066 |
| Clindamycin | 28 | 3,488 | 1,996 |
| Fusidic acid | 25 | 20,006 | 9,851 |
| Rifampin | 29 | 3,368 | 2,044 |
| Ciprofloxacin | 28 | 3,662 | 1,931 |

### Comparison of machine learning classifiers for AMR phenotypes

For all categories of classification, the hybrid method of using both the SNPs and genes as the predictors in one matrix, significantly underperformed compared to using just SNPs or just gene predictors, and is not reported here.

Classification using random forests always performed better when using the square root number of variables instead of the log_2_ number of variables. Therefore, only results from the random forest with a square root number of variables per tree are reported.

### Vancomycin

The vancomycin group contained a total of 11,066 SNP predictors. With the gene-centric approach, the pan-genome consisted of 4,865 genes.

Binary classification of vancomycin resistance revealed that genes are a superior predictor for this particular task. With the exception of the decision tree, Naïve Bayes, and AdaBoost, the gene-centric approach produced consistently superior f1 micro-scores with every algorithm. The highest scoring algorithm, the support vector machine, was able to achieve a mean 0.84 f1 micro-score (standard deviation 0.152) with gene predictors (Figure 3). Both the linear SVM and a radial basis kernel SVM performed similarly with large C parameters. In particular, the radial SVM dramatically improved in classification accuracy when increasing C from 0.1 to 10000 (Figure 4).

**Figure 4.**
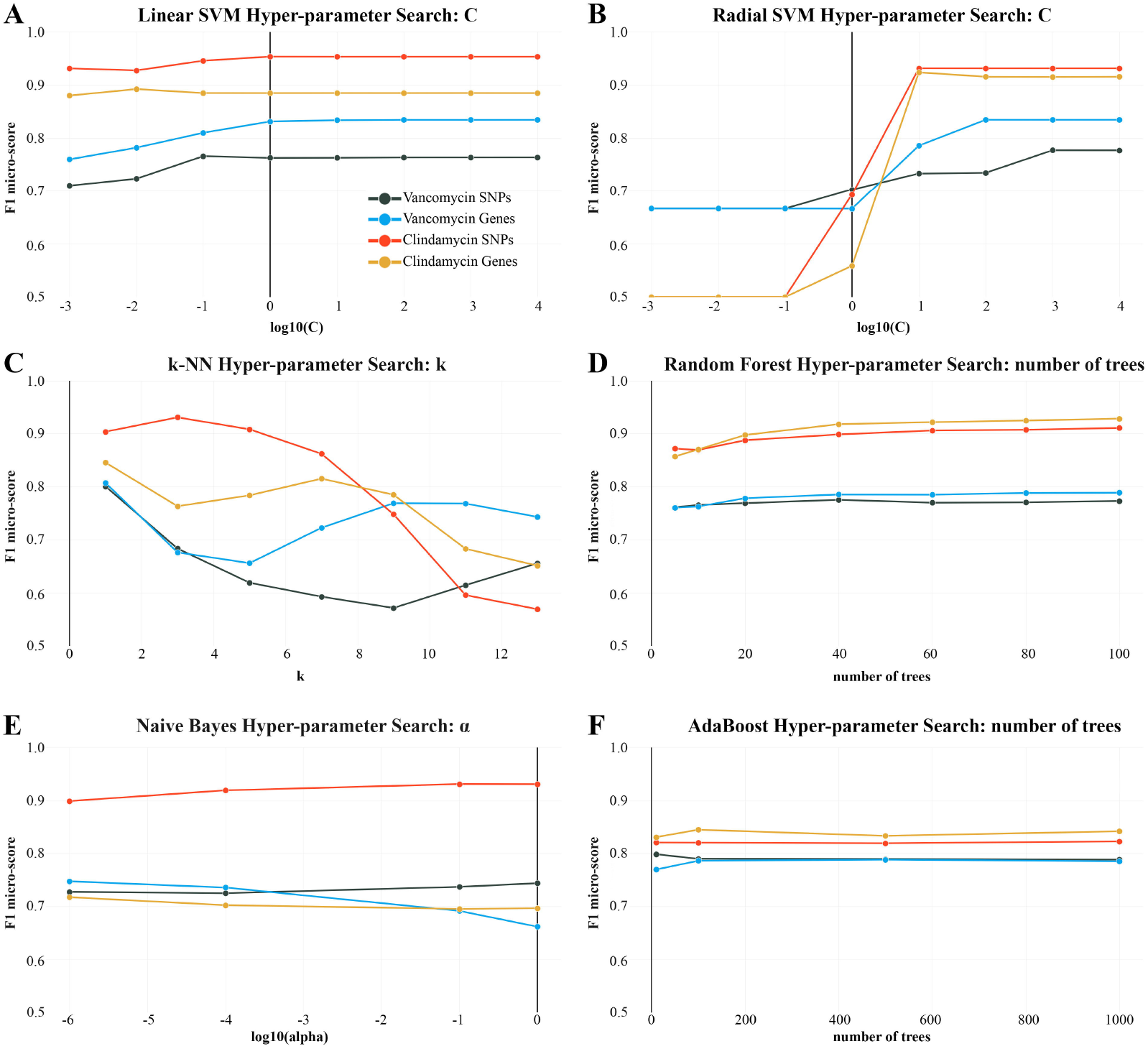
f1 micro-scores for each algorithm over the course of different hyper-parameter changes. k-NN (C.) and the SVM, with a radial basis function (B.) as a kernel, are most sensitive to hyper-parameter changes. Y-axis (f1 micro-score) is scaled from 0.5 to 1.0. The remaining plots (A. D. E. F.) show minimal changes to f1 micro-score from changing the respective hyper-parameter.

Outside of support vector methods, k-NN with k=1 performed with similar accuracy to the linear SVM, in the binary case, being very sensitive to the neighbor parameter: as shown in Figure 4, k-NN performed best when k=1, with subsequent degradation in accuracy. Conversely, greater consistency can be achieved with larger k values. For example, k=9 and k=11 reduce the total f1 micro-score compared to k=1, but have a smaller standard deviation when computing their mean f1 micro-score over many iterations (when k=1 std.=0.15 and when k=11 std.=0.11 for SNPs).

With SNP predictors, accurate binary AMR classification with a decision tree produced comparable results to gene-based SVMs and k-NN and required no hyper-parameter tuning. This method performed the best of all SNP predictor based methods.

With genes, ensemble based methods are less accurate than SVMs and neighbor methods in binary classification. Ensemble techniques did not show significant improvements when increasing the number of trees; although, adding more trees improved f1-scores slightly (Figure 4).

In the case of the multiclass classification, the core-SNP approach with a radial SVM performed the best, with an f1-score of 0.71 (Figure 3). k-NN (k=1) with genes performed roughly the same. Multiclass classification with tree based methods (random forest and decision tree) showed a 0.06 − 0.07 improvement to the f1 micro-score by using SNP variables.

### Clindamycin

The clindamycin group of 28 individuals contained an initial total of 1,996 SNP predictors. With the gene-centric approach, the pan-genome consisted of 6,949 genes.

Prediction of binary AMR classification with respect to clindamycin was most effective using SNPs (Figure 3). An f1 micro-score of 0.94 was achieved using a linear SVM and SNP predictors, compared to a lower micro-score of 0.89 using gene predictors. The radial SVM performed slightly worse than the linear variant and required a carefully chosen C parameter. As seen with other AMR classifications, larger C values were necessary for the radial SVM to perform adequately (Figure 4).

k-NN with SNPs produced an f1 micro-score of 0.92 with k=3. With gene predictors, this score decreased dramatically (0.85 score, k=1 was best with gene predictors). When using SNP predictors, any k > 3 marked a consistent degradation in algorithm prediction performance. A similar trend was seen in the variation of the f1-scores during cross-validation; k=3 yielded a lower standard deviation (std. =0.10) when compared to most other k values.

The random forest, with either genes or SNPs as predictors, performed competitively with SVM methods and neighbor methods in terms of f1 micro-score. The other ensemble method, AdaBoost, also generated similar performance between SNPs and genes as predictors but is significantly outperformed by random forests. Changing the number of estimators (trees) in either ensemble algorithm did not affect accuracy; although, having less than 20 trees in a random forest negatively impacted performance.

Using SNPs, and a large alpha value, Naïve Bayes achieved a 0.93 f1 score for clindamycin AMR prediction (Figure 3B). A large discrepancy between the gene predictor and SNP predictor methods was also found in Naïve Bayes AMR classification accuracy where using gene predictors resulted in a ~22% reduction in performance.

In the multiclass AMR classification, SVMs (with SNPs) again outperformed all other algorithms with a 0.71 f1 micro-score (for both kernels). Using the other described algorithms results in lower, and generally uniform, f1 scores, except for AdaBoost which underperforms with a score of 0.55 using both SNPs and genes.

### Fusidic acid

The fusidic acid group contained a total of 9,851 SNP predictors. With the gene-centric approach, the pan-genome consisted of 4,727 genes used as predictors.

Multiclass classification of the fusidic acid group, with four possible AMR categories, showed that gene variables dominated as the best predictor across all tested algorithms with an average f1 score improvement of 0.06. Gene-centric k-NN performed the best, with a mean f1 micro-score of 0.70 and a standard deviation of 0.17 (Figure 3E). Like with the other phenotypes, SVMs (linear and radial) were top performers, achieving a 0.67 score. In the case of ensemble methods, using genes, both the decision tree and AdaBoost were out-performed by random forests, using 100 trees, which achieved an almost comparable score to SVMs and k-NN (Figure 3E).

### Rifampin

With a total of 29 individuals in the rifampin group, a total of 2,044 core SNPs were generated. The pan-genome consisted of 7,805 unique genes.

Multiclass prediction of rifampin resistance levels produced the most successful results among the multiclass tests, with a mean f1 micro-score 0.75 (0.173) with a decision tree and SNPs. Radial kernel-based SVMs and random forests provided similar accuracy (Figure 3F), SNP predictors produced superior accuracy with all algorithms except for k-NN and Naïve Bayes. The decision tree showed the largest discrepancy when changing predictor type, going from top classifier with SNPs to nearly the lowest scoring classifier with genes, showing extreme sensitivity to predictor type.

### Ciprofloxacin

The ciprofloxacin group contained 28 individuals. The core-SNPs were 1,931 in total. The total number of pan-genes was 6,975.

The results of ciprofloxacin multiclass prediction proved inferior to other phenotypes, despite the uniformity of the training and test set distribution. Although the decision tree with SNPs produced the top f1 micro-score of 0.54 (Figure 3G), it also produced the largest standard deviation (0.20) in its f1 micro-score. Alternatively, Naïve Bayes with SNPs produced a slightly lower mean score of 0.52, but with greater model stability (standard deviation of 0.15).

## Discussion

The rapid evolvability of AMR systems and the subsequent surge of extended-spectrum resistance phenotypes (Shaikh et al., 2015) has dramatically impacted the characterization of virulent microbes. In this manuscript, we inspected the predictability of AMR using six ML algorithms, and the results suggest a promising ML-based approach for the prediction of binary AMR classification (i.e. resistant versus susceptible). The employment of multiple different learning algorithms on a small set of in-house samples, combined with “cloud knowledge”, revealed that support vector machines, k-NN, and random forests can be trained to high accuracy with less than 35 samples, using thousands or tens of thousands of predictors (Table 2). Using sequence data, two types of biological predictors are immediately available: core-genome SNPs and gene presence/absence matrices, both of which yield similar levels of prediction accuracy. However, based on our results, one set of predictors may prove particularly effective at AMR prediction than other predictor sets for a particular phenotype. For example, SNPs seem to be the preferred predictors for clindamycin predictive tasks (Figure 3B), whereas vancomycin favors gene predictors (Figure 3A). Limited sample sizes continue to make multiclass AMR challenging, as demonstrated in the cases of fusidic acid and ciprofloxacin (Figure 3E to Figure 3G).

Susceptibility testing of *Elizabethkingia* strains investigated in this study revealed some species-specific trends with regards to MICs and MBCs (Supplementary Table 1). For instance, with the exception of strain R26, the *E. anophelis* clindamycin MICs (all 1 mg/L) and MBCs (1-8 mg/L) were consistently higher than all other species investigated. The *E. meningoseptica* MICs and MBCs for vancomycin were higher than most other strains investigated. When investigating the genetics underlying AMR, annotation-based discovery can only be effective when genetic annotation for the genotype responsible for the observable AMR is available and when homology exists. A successful case is seen in a recent study (Bosse et al., 2017) that concluded a 100% correlation between annotated putative genes and targeted AMR phenotypes. In our case, annotation of both genomes disclosed a small number of genetic components associated with the corresponding MIC results. For example, *vanA* and *vanH*, genes belonging to the *vanA* operon, were identified in the extremely vancomycin resistant *Enterococcus faecium* strain 805447/07 (MIC value of 256 mg/L, Supplementary Table 1). The *vanA* operon is a genetic element that provides resistance to vancomycin by facilitating the replacement of a dipeptide in peptidoglycan synthesis, making vancomycin less likely to bind to peptidoglycan precursors and inhibit cell wall synthesis (Perichon and Courvalin, 2009).

Annotation-based AMR detection can be complicated by the accumulation of mutations at these gene sites, reducing the effectiveness of alignment-based homology searches using a database (Rost, 1999). Similarly, situations where the resistance phenotype is the product of genetic pathways with undescribed genes, or with genes of an unknown function, can make meaningful annotation-based conclusions challenging. For example, every *Elizabethkingia* strain was annotated as the accessory gene *vanW*, believed to play a part in vancomycin resistance but with unknown function (Guardabassi et al., 2005;Raygoza Garay et al., 2016). Moreover, there is no evidence that *vanW* plays any role in the defense mechanism of *Elizabethkingia*, and this gram-negative genus is likely intrinsically resistant to vancomycin due to the outer membrane’s impermeability to large glycopeptides. However, these annotations failed to provide satisfactory support to the diverse MIC values observed in our *Elizabethkingia* strains (Supplementary Table 1 and Supplementary Table 2).

In the post-NGS era, whole-genome sequencing (WGS) provides a convenient means to explore genomic variants of entire chromosomes. Raw genomic data is often represented as genome assemblies or sequenced reads and has historically been difficult to extract genotype-phenotype relationships from. Producing genome-wide predictors, like SNPs, has become a conventional approach, but the SNPs generated by NGS techniques can be prone to noise (Briskine and Shimizu, 2017;Wu et al., 2017); this, in addition to limited sample sizes and the large quantity of SNP predictors, has imposed significant challenges for many statistical modeling approaches (Lange et al., 2014). This phenomenon is known as the ‘curse of dimensionality’ (Bellman, 2010). The dimensionality issues, in combination with noisy predictors, can lead to problems of overfitting and model mis-identification (Sun et al., 2019). Careful consideration of the genomic data, with respect to its usage in algorithms, must be applied for appropriate predictive modeling.

Prediction accuracy is data-dependent and relies on the quality of the predictor variables used as input. Also, some algorithms, like SVMs and k-NN, require careful hyper-parameter tuning to function properly (Figure 4). In this study, we tested different biological predictor types and found that, depending on the AMR phenotype, either genes or SNPs reliably produce better results. For example, the multiclass fusidic acid classification greatly favors genes over SNPs, while the vancomycin multiclass classification performs better using SNP predictors. Unlike the core-SNPs, our gene-centric model can produce predictors from genomic regions outside of the core-genome, and gene presence/absence predictors do not suffer from small sequencing errors due to their reliance on alignments, which can tolerate some errors using score matrices. This is also advantageous when the strain exhibits an AMR phenotype that is based around a gene product, such as fusB-type fusidic acid resistance. In these genic AMR cases, the effect of mutational changes in the core genome might be too insignificant to be reliably explanatory for AMR phenotypes.

A common problem seen when analyzing biological data from individuals that are not part of a controlled population is the relative lack of available samples compared to the wealth of genomic data. The curse of dimensionality characterizes this problem as requiring a tremendous amount of sample data to guarantee that each potential combination of SNP/gene predictor exists within the dataset. As shown in Table 2, the number of variables can far outnumber the sample size and, the ratio of samples to SNPs can be as low as 0.002 (Table 2), depending on the core genome. Such small sample sets are challenging to model using traditional statistical methods, often requiring feature selection or regularization, as shown in (Wu et al., 2009). For example, in a 2018 study (Manavalan et al., 2018) this problem was approached using random forests to reduce the total number of variables assigned to each tree, and resulted in an improved 87% accuracy, using leave-one-out cross-validation. Regularization is a core technique in machine learning used to mitigate overfitting. The SVM, for instance, can allow for a wider hyperplane when optimizing, permitting the misclassification of certain training samples in exchange for higher generalizability. The use of support vectors also helps alleviate issues with small sample size, because the decision boundary can be derived from a small subset of the training data. Our results in Figure 3 demonstrate the merit of SVM-based classifiers with a small sample size. The high classification accuracy was also achieved in part due to the inbuilt regularization of determining the margins of the decision boundary, since margin generation is independent of the features’ dimensionality. The benefits from SVM regularization are maintained even with a large sample size (Araya and Hazelhurst, 2009). Recent studies have shown success with ML in predicting biofilm inhibiting peptides (Gupta et al., 2016), identifying bacteriophage virion proteins (Manavalan et al., 2018) and productivity estimates using microbiome composition (Chang et al., 2017). Through its effective modeling of the relationships between predictors and reduction of the effects of noisy data, ML has also made a significant impact on genomics, where it has been used for expression prediction and genomic element recognition (Libbrecht and Noble, 2015).

Together with this research, we suggest that this sample size dilemma can be significantly lessened by capitalizing on the wealth of information stored on cloud services. There are currently more than 150,000 prokaryotic genome assemblies available on the NCBI. We used this community-driven “cloud knowledge” to increase our sample population by 57% and 33% for the vancomycin and clindamycin groups respectively (Table 2), compared to just using in-house *Elizabethkingia* strains and also produced a more uniform phenotypic distribution which made prediction more feasible.

Decreasing sequencing costs and the subsequent increase in availability of genomic information stored on cloud services like NCBI and the European Nucleotide Archive means that “cloud knowledge” will be an effective means of improving sample sets for AMR prediction, provided proper phenotyping is performed. Practical use of a computational AMR prediction pipeline in a clinical or hospital setting will also depend on a computationally efficient predictor generation method. Also, given the large, diverse collection on the NCBI, constructing a core-genome could only be advantageous for species/strains that share common ancestry; the same approach can be problematic when including unrelated genera, owing to the decrease in conserved genomic regions, reduced numbers of predictors and possible losses of functional homology. Further, multiple-sequence alignment software must be able to align 1000’s of genomes in a time-efficient manner. Recently, alignment-free variant calling has become an effective alternative and does not demonstrate the excessive time-complexity of traditional methods (Zielezinski et al., 2017). Our results also show that gene presence matrices are equally as effective as SNPs, sometimes better, in most cases (Figure 3). With current-generation, high-speed annotation software, multiple-sequence alignment may be unnecessary, and emerging pathogens with unknown AMR resistance can rapidly and easily be annotated and predicted. Supported by the latest improvements in metagenomic assembly (Olson et al., 2017), it is feasible to directly sequence from infected tissue and produce assemblies from microbial communities for use in prediction. To conclude, we expect ML predictive pipelines, in combination with metagenomics, will shift the paradigm from phenotype-based diagnostics to data-driven prediction for AMR detection and outbreak prevention.

## Acknowledgements

This research is also supported by the NSF-MRI 1626257 for P.H. and C.C.; the work presented in this report reflects the support from the USDA HATCH project OKL03011 of C.C.. Data analysis was completed with support from the High Performance Computing Center Facilities at Oklahoma State University.

